# Time-resolved hemodynamic responses to sentence-level speech perception, production, and self-monitoring

**DOI:** 10.64898/2026.04.13.715885

**Authors:** Teng Ieng Leong, Aotong Li, Jian Hwee Ang, Barry Lee Reynolds, Cheok Teng Leong, Chi Un Choi, Martin I. Sereno, Defeng Li, Victoria Lai Cheng Lei, Ruey-Song Huang

## Abstract

Functional magnetic resonance imaging (fMRI) has been widely utilized to explore the neural mechanisms underlying speech processing. However, the intertwining of perception and production that exists in real-world scenarios remains underexplored due to challenges such as gradient noise and head motion artifacts from speaking. Previous research has often employed sparse-sampling designs, pausing image acquisition intermittently to present auditory stimuli or record overt speech. While this approach mitigates some challenges, it cannot capture continuous brain activity during speech processing and does not separate the mixed hemodynamic responses to external and self-generated speech occurring in succession. We overcame these limitations and continuously scanned thirty-one participants as they listened to and recited English sentences. Through independent component analysis (ICA), we decomposed each functional scan into spatially independent components (ICs), identifying task-related ICs in the superior temporal cortex, inferior frontal gyrus, and orofacial sensorimotor cortex. These ICs demonstrated time-resolved hemodynamic responses corresponding to distinct stages of speech perception, planning, and production. A linear subtraction between the IC time courses from the listening-reciting (perception-to-production) and listening (perception-only) tasks further revealed a secondary hemodynamic response to self-generated speech in the superior temporal cortex. Furthermore, we established precise temporal relationships between overt speech output and the peak, rise, and fall of hemodynamic responses for each independent component. Together, we present a methodological framework that can inform future fMRI studies on naturalistic tasks involving the perception of external auditory stimuli and monitoring of self-generated sounds.

## 1. Introduction

The ability to distinguish between temporal activations in the brain regions associated with auditory perception and self-monitoring is vital for advancing our understanding of cognitive processes across various domains, particularly in language and music processing. Despite significant progress in mapping the cortical representations of language processing with functional magnetic resonance imaging (fMRI) (Price, 2012), the temporal dynamics of brain activity during naturalistic sentence-level speech perception and production have not been sufficiently investigated (Janssen & Mendieta, 2020; Matchin et al., 2019). Investigating these processes using fMRI presents inherent challenges, notably the masking effects of gradient noise on auditory inputs and complications arising from head motion during overt speech production (Gracco et al., 2005; Jolly et al., 2020). To address these challenges, researchers have used sparse-sampling fMRI designs to avoid the interference of scanner noise on speech perception and production (Blackman & Hall, 2011; Perrachione & Ghosh, 2013). In a sparse-sampling fMRI experiment, image acquisition is intermittently paused to allow auditory stimulus presentation or the recording of overt speech against a silent background (Merrett et al., 2021), which also avoids recording head motion artifacts while speaking. However, optimized sparse-sampling fMRI designs depend on estimating precise time windows to record the maximum brain activity following stimulus presentation or speech output. Determining the exact interval between stimulus onset and the peak hemodynamic response is challenging for paradigms involving sentence processing, as the hemodynamic response can begin to rise while the stimulus is still being presented.

Although the interleaved silent steady sparse-sampling method attempts to mitigate discrepancies due to varying stimulus durations (Schwarzbauer et al., 2006), it still falls short of capturing the complete temporal profile of the hemodynamic response elicited by sentence-level stimuli (Wise & Geranmayeh, 2016). In our recent studies, we overcame the challenges associated with scanner noise and head motion, achieving continuous fMRI sampling with real-time auditory feedback during naturalistic speech perception and production tasks (Lei et al., 2024; Lei et al., 2025). Our studies demonstrated that continuous sampling could resolve the phases of periodic hemodynamic fluctuations, enabling further exploration of complex spatiotemporal brain dynamics within and across regions, which are inaccessible with sparse-sampling designs. However, a persistent challenge for continuous sampling is disentangling the overlapping blood-oxygenation-level–dependent (BOLD) signals elicited by external and self-generated speech occurring in close temporal proximity. While typical experimental designs for studying the neural basis of speech processing usually consist of isolated stimuli and responses (Vigneau et al., 2006; Walenski et al., 2019), real-world communication often involves overlapping or sequential intervals of speech perception and production. For example, an interpreter actively listens to a spoken sentence in a source language and renders it into a target language, either simultaneously or consecutively (Paradis, 1994; Hervais-Adelman et al., 2015). In a similar vein, collaborative singing serves as a prime example of music processing that involves both external auditory input and self-monitoring, where vocalists listen to other vocalists or instrumental accompaniment while simultaneously monitoring their own vocal output to achieve harmony and synchronize their performance. Both scenarios highlight the auditory system’s capacity to process both external stimuli and self-generated sounds, which leads to temporally mixed hemodynamic responses in the auditory regions.

In this study, subjects were scanned continuously while they engaged in two tasks: passively listening to spoken sentences, and actively listening to, memorizing, and reciting spoken sentences. Using independent component analysis (ICA) (Beckmann & Smith, 2004; Davis et al., 2014; Janssen & Mendieta, 2020; McKeown et al., 1998; Plante et al., 2017), our aim is to decompose and identify independent brain processes associated with speech perception, planning, production, and self-monitoring. By addressing the challenges of overlapping BOLD signals, this research seeks to unravel the full temporal profiles of hemodynamic responses during complex speech tasks, contributing to a deeper understanding of auditory processing in real-world communication.

## 2. Methods

### 2.1. Participants

The study analyzed fMRI datasets of 31 subjects (22 females, 9 males; mean age: 23.3 ± 4 years) recruited by the Brain and Language Project at the University of Macau (Lei et al., 2024). All subjects were native Chinese (Mandarin) speakers who acquired English as a second language between the ages of 4 and 13 (mean age of acquisition = 7.1 ± 2.1 years). English proficiency was assessed based on their most recent standardized test results (an IELTS score of 6.5 or above, or an equivalent qualification). All subjects gave written informed consent in accordance with the protocols approved by the Research Ethics Committee of the University of Macau. All subjects had normal or corrected-to-normal vision and reported no history of neurological disorders.

### 2.2 Experimental procedures and setup

To minimize head motion during overt speech production, we made an individualized facial mask for each subject using thermoplastic sheets (1.6 mm H-board, Sun Medical Products Co., Ltd.). All subjects then wore their masks and completed a training session to practice speaking while maintaining head stillness in a mock scanner (Shenzhen Sinorad Medical Electronics Co., Ltd.) (Fig. 1). During the training, real-time head motion was monitored using a sensor attached to their forehead (MoTrak, Psychology Software Tools, Inc.). Subjects would hear a warning sound (“ding”) if their head motion exceeded 1 mm in translation or 1° in rotation, allowing them to monitor their head position while speaking.

**Fig. 1.**
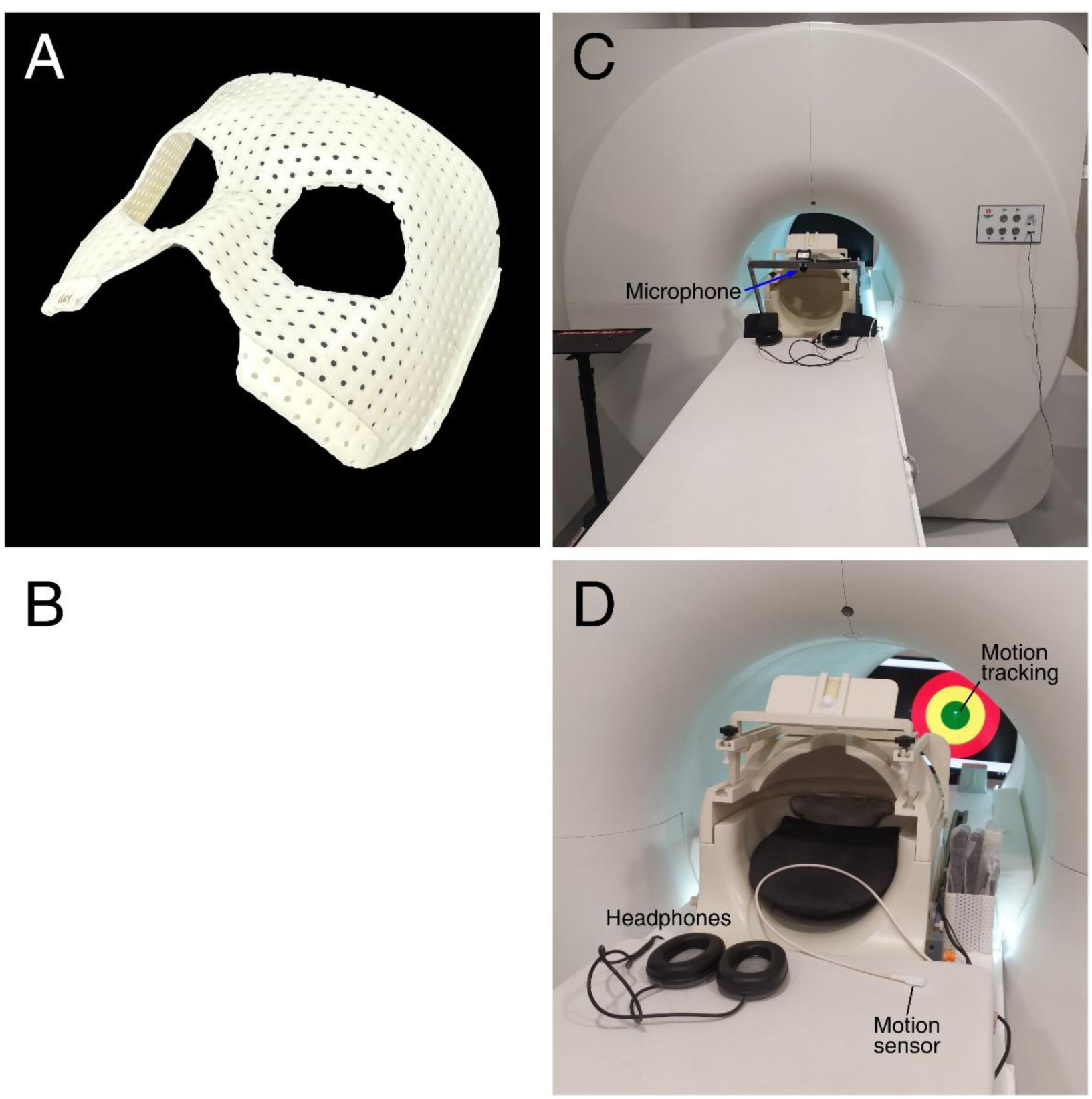
Experimental setup for speech perception and production tasks. (A) A custom-molded mask with “The phantom of the opera” style for head immobilization. (B) Demonstration of subject positioning in a Siemens 32-channel head coil. The subject wears a mask and a pair of MR-compatible headphones, with a microphone placed in front of her mouth. The gaps surrounding the mask and headphones are filled with resin clay and silicone gel. (C) An MRI simulator (bore diameter = 60 cm) and a mock coil equipped with a microphone and headphones. (D) A close-up view of the mock coil. The motion sensor is affixed to the subject’s forehead during simulator training.

Prior to an fMRI session, subjects would wear a mask and earplugs, with their ears fully covered by a set of MR-compatible headphones (OptoACTIVE II, OptoAcoustics Ltd.), and then lay supine with their heads supported by resin clay in a head coil. The mask and headphones were further stabilized by filling resin clay and silicone gel in the gap between the head and the inner wall of the head coil, which also effectively attenuated scanner noise.

To ensure clear and intelligible delivery of auditory stimuli during fMRI experiments, a dedicated noise cancelling system was implemented. The MR-compatible headphones and microphone (FORMRI-III, OptoAcoustics Ltd.) were connected to a real-time active noise cancellation system via fiber optic cables passing through a waveguide connecting the scanner room and control room. This system analyzed the power spectrum of gradient noise of an echo planar imaging (EPI) pulse sequence and actively cancelled the noise, effectively suppressing scanner-generated interference during speech perception and production tasks.

In addition to active noise cancellation, passive attenuation was achieved through the combined use of earplugs, resin clay, and an acoustic foam (4.5 cm thick) lining the scanner bore. These measures substantially reduced ambient gradient noise. All subjects reported no difficulty in hearing or understanding the auditory stimuli, including external and self-generated speech, throughout each functional scan. Together, these integrated measures for countering head motion and scanner noise enabled continuous functional imaging during sentence-level speech perception and overt reciting tasks.

### 2.3. Experimental design

All subjects participated in twelve scans of six sentence-level perception and production tasks for both Chinese and English sessions (Lei et al., 2024). Of the twelve scans comprising six language tasks, the present study focused on listening and listening-reciting tasks from the English session. Each task was performed in two 256-s functional scans with a rapid phase-encoded design (Chen et al., 2019; Lei et al., 2024), each consisting of sixteen 16-s consecutive trials, i.e., 16 cycles/scan. Within each trial of the listening task, subjects passively listened to a spoken sentence for 5 s (perception phase: 0–5 s), followed by 11 s of rest (5–16 s). Within each trial of the listening-reciting task, subjects passively listened to and memorized a spoken sentence for 5 s (perception phase: 0–5 s), followed by 5 s of reciting from memory (production phase: 5–10 s), and 6 s of rest (10–16 s). Visual cues, including an ear icon for the perception phase and a mouth icon for the production phase, were presented at the center of a rear-view LCD screen. All stimuli were systematically constructed with 15–17 syllables per sentence, following the subject-verb-object structure, e.g., “She has posted photographs of her friends on bulletin boards.” All stimuli were built from high-frequency vocabulary sourced from the *Longman Communication 3000 list* (https://www.lextutor.ca/freq/lists_download/longman_3000_list.pdf) and the *Collins Dictionary English* (https://www.collinsdictionary.com/dictionary/english/corpus), and validated for linguistic appropriateness by a professor of linguistics.

### 2.4. Image acquisition

All images were acquired with a 32-channel head coil in a 3T MRI scanner (Siemens MAGNETOM Prisma) at the Centre for Cognitive and Brain Sciences, University of Macau. For functional imaging, whole-brain functional images were acquired using a blipped-CAIPIRINHA simultaneous multi-slice (SMS), single-shot EPI sequence with the following parameters: acceleration factor = 5; interleaved ascending slice order; repetition time (TR) = 1000 ms; echo time (TE) = 30 ms; flip angle = 60°; 55 axial slices; field of view (FOV) = 192×192 mm²; matrix size = 64×64; voxel size = 3×3×3 mm³; bandwidth = 2368 Hz/Px. Each functional scan comprised 256 volumes (excluding 6 dummy volumes), resulting in an effective scan time of 256 seconds per scan. Additionally, two scans of T1-weighted structural images were acquired, matching the slice center and orientation of functional scans, using an MPRAGE sequence (TR = 2300 ms; TE = 2.26 ms; inversion time (TI) = 900 ms; flip angle = 8°; 256 axial slices; FOV = 256×256 mm²; matrix = 256×256; voxel size = 1×1×1 mm³; bandwidth = 200 Hz/Px; scan time = 234 s).

### 2.5. Image preprocessing

All raw images (DICOM .ima files) were converted to Analysis of Functional NeuroImages (AFNI; https://afni.nimh.nih.gov) .BRIK files using the AFNI *to3d* command. Functional images were motion-corrected by registering the volumes in all scans to the first volume of the seventh scan (immediately after a structural scan) using the AFNI *3dvolreg* command, yielding motion parameters in six degrees of freedom for each scan (Fig. 2). Subsequently, the motion-corrected functional images were coregistered to the structural images using the *csurf* package (https://pages.ucsd.edu/~msereno/csurf) (Sereno et al., 2022). For each subject, two sets of T1-weighted structural images were aligned and averaged prior to cortical surface reconstruction using FreeSurfer 7.2 (https://surfer.nmr.mgh.harvard.edu) (Dale, 1999; Fischl et al., 1999a).

**Fig. 2.**
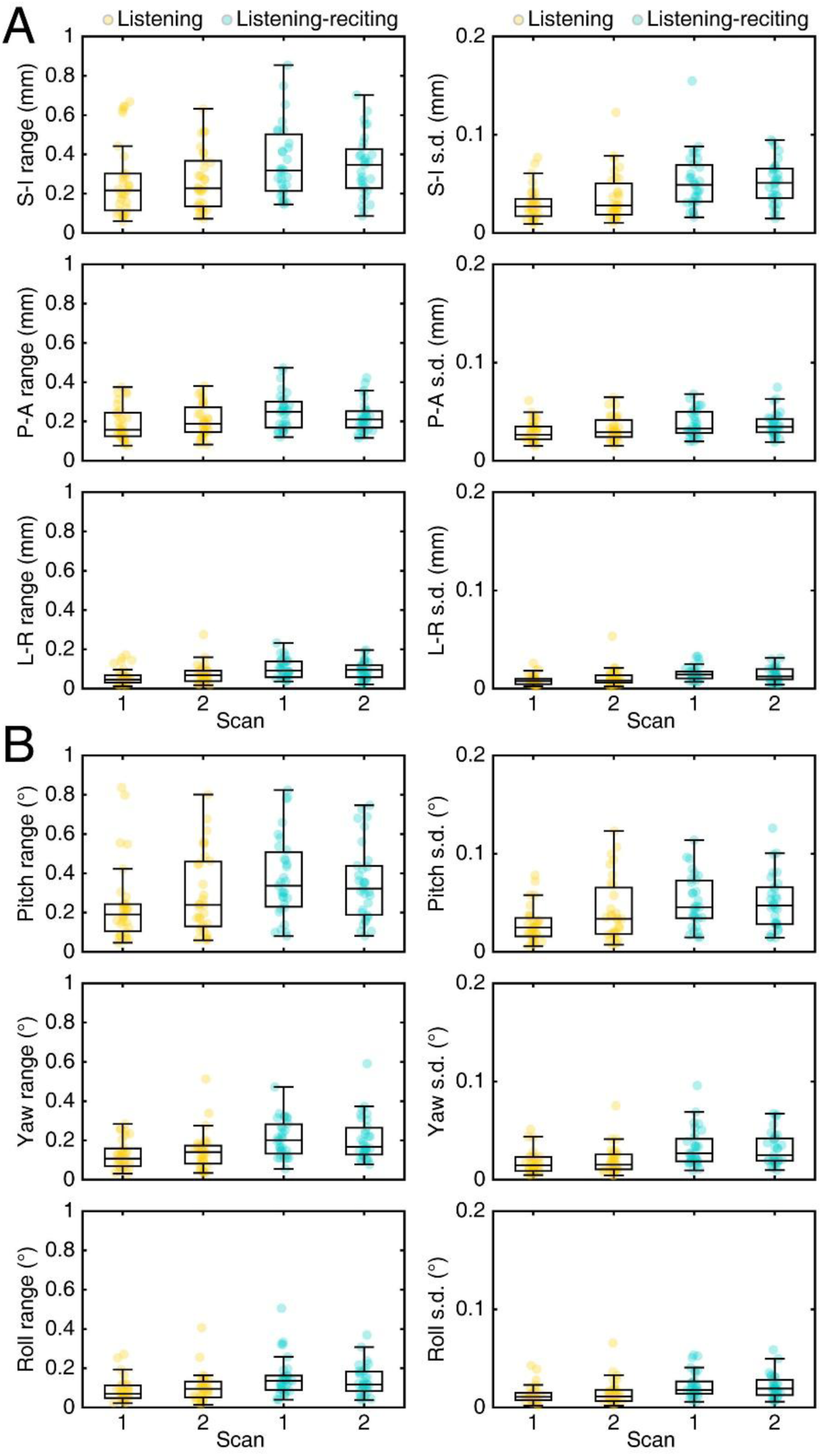
Distribution of motion parameters in six degrees of freedom. (A) Head displacement in the superior-inferior (S-I), posterior-anterior (P-A), and left-right (L-R) directions. (B) Head rotation in pitch, yaw, and roll. Left column: distribution of the range (maximum - minimum) of motion within each scan across subjects. Right column: distribution of the standard deviation (s.d.) of motion parameters within each scan across subjects. Each color marker indicates the data point from a single subject (*N* = 31).

### 2.6 Audio analysis

The soundtracks of auditory stimuli, speech output, and Transistor-Transistor Logic (TTL) signals in sync with the EPI pulse sequence were recorded simultaneously using the OptiMRI 3.1 software (OptoAcoustics Ltd.). The timings of onset and offset of speech production in the listening-reciting task were manually identified from the speech-output channel using the Audacity software (https://www.audacityteam.org).

### 2.7 Independent component analysis

Following motion correction and registration, the 4D dataset (64×64×55×256 data points) of each functional scan from individual subjects was subjected to probabilistic independent component analysis (Beckmann & Smith, 2004) using the FSL MELODIC toolbox (Version 3.15; https://fsl.fmrib.ox.ac.uk/fsl/docs/resting_state/melodic.html). The analysis resulted in a spatial map (64×64×55 points of Z-statistics values) and a time course (256 points) for each independent component (IC) decomposed from each functional scan of each subject (Figs. 3-6). Each IC spatial map (voxel-based) was displayed on the cortical surface of each subject using the FreeSurfer *paint* command, yielding a paint file (.w) consisting of a Z-statistics value at each vertex on the cortical surface. This subject-level surface-based IC spatial map was morphed and resampled to the cortical surface of the template brain *fsaverage* using the FreeSurfer *mri_surf2surf* command (https://freesurfer.net/fswiki/mri_surf2surf). We then identified and averaged ICs with similar spatial maps across subjects over the cortical surface of *fsaverage*. Surface-based group-average maps reveal consistent activations that are spatially aligned on a vertex-by-vertex basis (Fischl et al., 1999b; Sereno et al., 2022; Sood & Sereno, 2016). Misalignment or a random mix of positive and negative values at the same location would result in weak to no activations in the group average map. Each surface-based IC spatial map was overlaid with borders of regions of interest (ROIs) based on the HCP-MMP1 atlas for precise localization and interpretation of activations (Glasser et al., 2016).

**Fig. 3.**
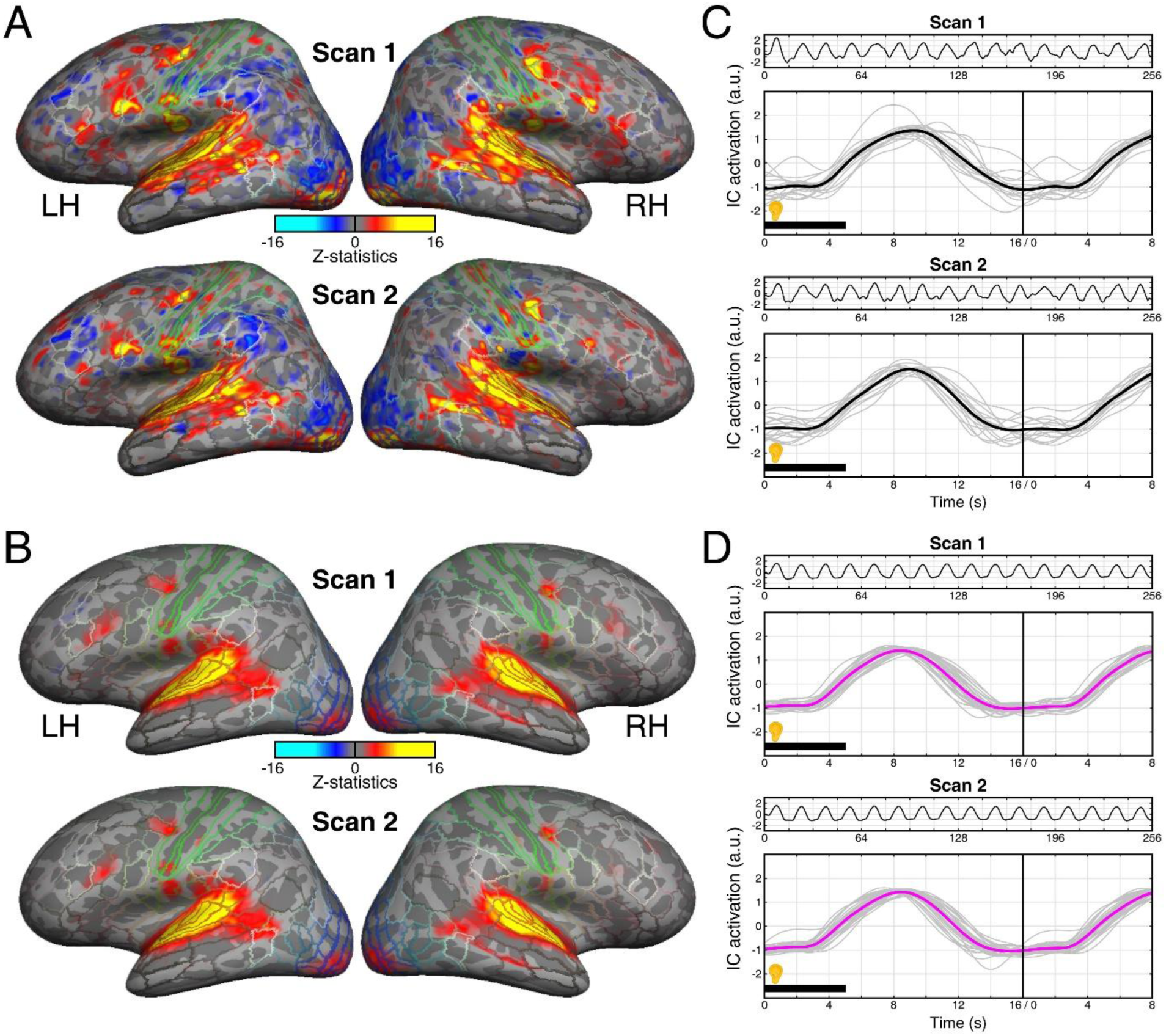
Spatial maps and time courses of the STC IC in the listening task. (A) Spatial maps of the STC IC from two scans of a single subject. LH: left hemisphere; RH: right hemisphere. The maps are overlaid with the borders of regions of interest (ROIs) defined in the HCP-MMP1 atlas (Glasser et al., 2016). (B) Group-average (*N* = 31) spatial maps of the STC IC from two scans. (C) Time courses associated with the STC IC in (A). The first and third panels show the full 256-s time courses for both scans. The second and fourth panels show the time courses in 24-s epochs (gray curves) and their averages (black curves) within each scan. (D) Time courses associated with the STC IC in (B). The first and third panels show the full 256-s time courses averaged across 31 subjects for each scan. The second and fourth panels show the time courses in 24-s epochs averaged within each subject (gray curves) and across 31 subjects (magenta curves) for each scan. a.u.: arbitrary unit.

**Fig. 4.**
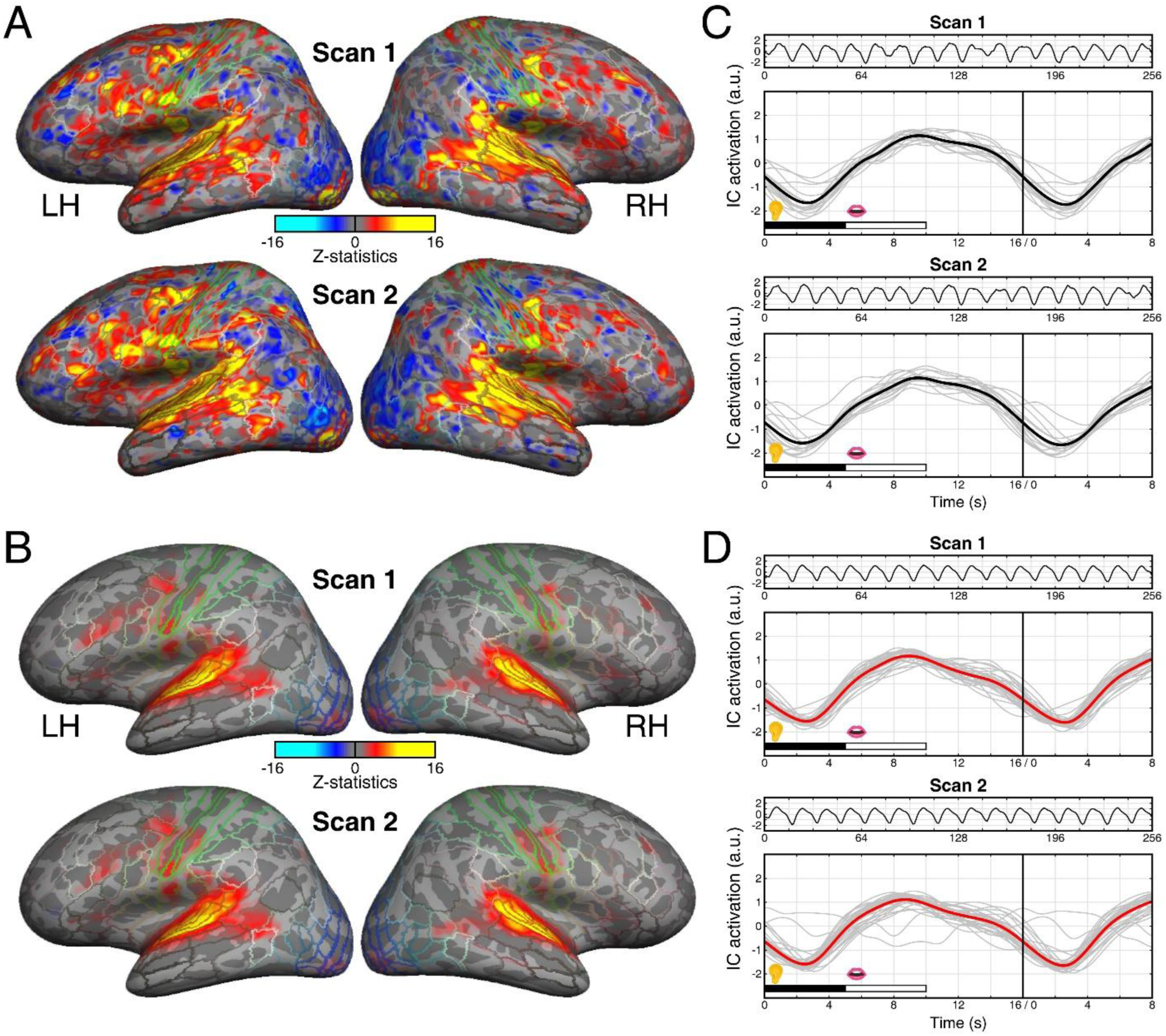
Spatial maps and time courses of the STC IC in the listening-reciting task. (A) Spatial maps of the STC IC from two scans of a single subject. (B) Group-average (*N* = 31) spatial maps of the STC IC from two scans. (C) Time courses associated with the STC IC in (A). The first and third panels show the full 256-s time courses for both scans. The second and fourth panels show the time courses in 24-s epochs (gray curves) and their averages (black curves) within each scan. (D) Time courses associated with the STC IC in (B). The first and third panels show the full 256-s time courses averaged across 31 and 29 subjects for Scan 1 and Scan 2, respectively. The irregular IC time courses from two subjects in Scan 2 were excluded from group averaging and subsequent analyses. The second and fourth panels show the time courses in 24-s epochs averaged within each subject (gray curves; *N* = 31 for Scans 1 and 2) and across subjects (red curves; *N* = 31 for Scan 1 and *N* = 29 for Scan 2). All conventions are consistent with those in Fig. 3.

**Fig. 5.**
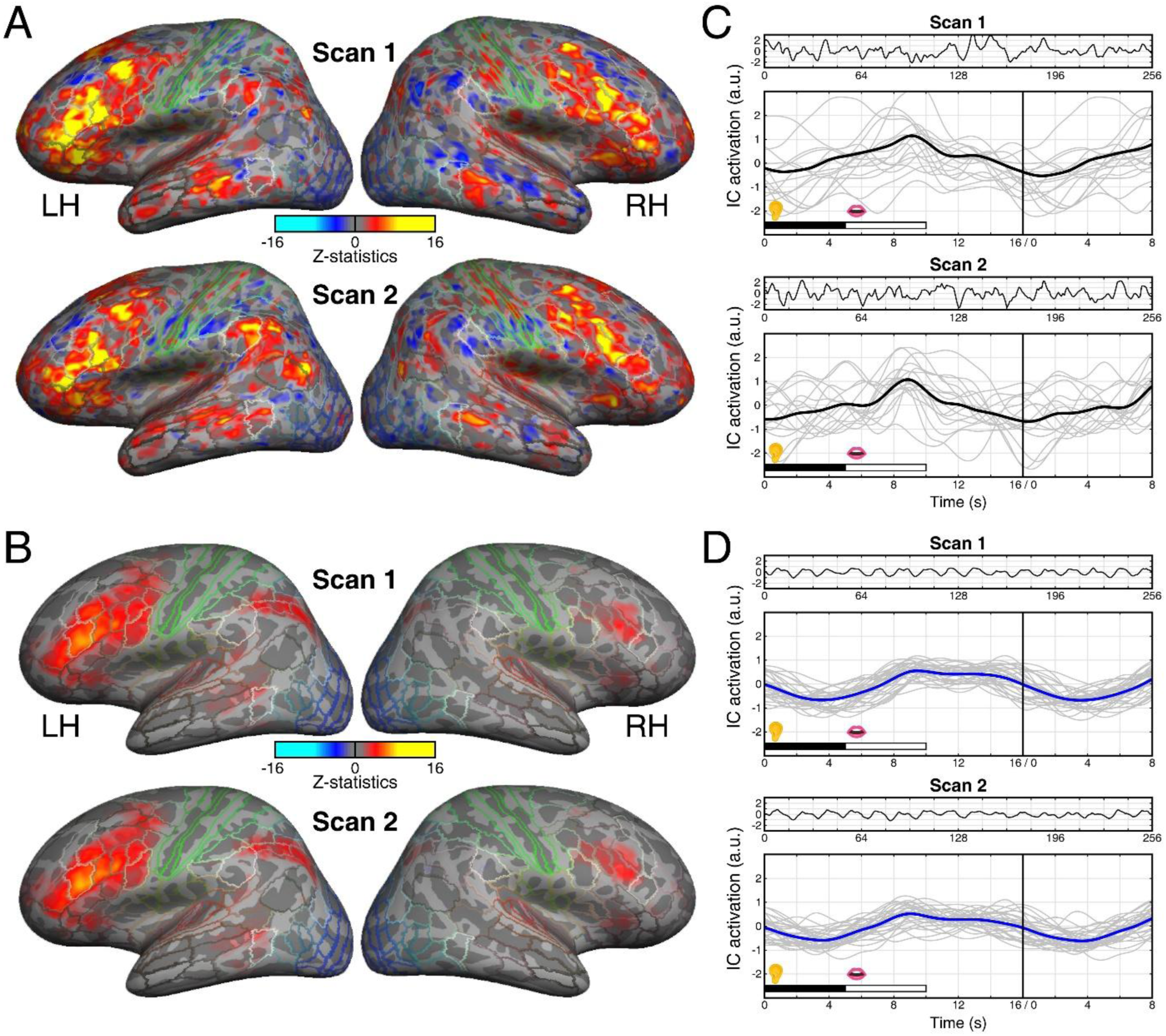
Spatial maps and time courses of the IFG IC in the listening-reciting task. (A) Spatial maps of the IFG IC from two scans of a single subject. (B) Group-average (*N* = 31) spatial maps of the IFG IC from two scans. (C) Time courses associated with the IFG IC in (A). The first and third panels show the full 256-s time courses for both scans. The second and fourth panels show the time courses in 24-s epochs (gray curves) and their averages (black curves) within each scan. (D) Time courses associated with the IFG IC in (B). The first and third panels show the full 256-s time courses averaged across 31 and 29 subjects for Scan 1 and Scan 2, respectively. The IC time courses from two subjects in Scan 2 (the same as those excluded in Fig. 4) were excluded from group averaging and subsequent analysis. The second and fourth panels show the time courses in 24-s epochs averaged within each subject (gray curves; *N* = 31 for Scans 1 and 2) and across subjects (red curves; *N* = 31 for Scan 1 and *N* = 29 for Scan 2). All conventions are consistent with those in Fig. 3.

For each subject *S*, an IC time course *X*(*t*), *t* ∈ {1, 2, …, 256} s, was upsampled 100 times to *X’*(*t’*), *t’* ∈ {0, 0.01, 0.02, …, 256} s, using the cubic spline interpolation algorithm implemented in the Matlab *interp1* function. Note that the data point at *t* = 0 was duplicated from that at *t* = 1 prior to the interpolation. The upsampled time course was then segmented into 16 overlapping 24-s epochs, 𝑋^′^*_E_* (𝑡^′^*_e_*), *E* ∈ {1, 2, …, 16} and 𝑡^′^*_e_* ∈ {0, 0.01, 0.02, …, 24} s. Each epoch began at the onset (0 s) of a 16-s trial and continued for 8 s into the subsequent trial, ensuring that the complete hemodynamic response profile was included (panels C and D in Figs. 3-6). The data points in the 16 to 24-s range of the last (16th) epoch were obtained by averaging the corresponding data points from the first 15 epochs. For each IC time course, sixteen epochs 𝑋^′^*_E_* (𝑡^′^*_e_*) were averaged within each subject, resulting in 𝑋^′^*_S_*(𝑡^′^*_e_*); a group-average epoch 𝑋^′^*_G_* (𝑡^′^*e*) was obtained by further averaging epochs 𝑋^′^*_S_* (𝑡^′^*_e_*), *S* ∈ {1, 2, …, 31}, across subjects (Fig. 7).

**Fig. 6.**
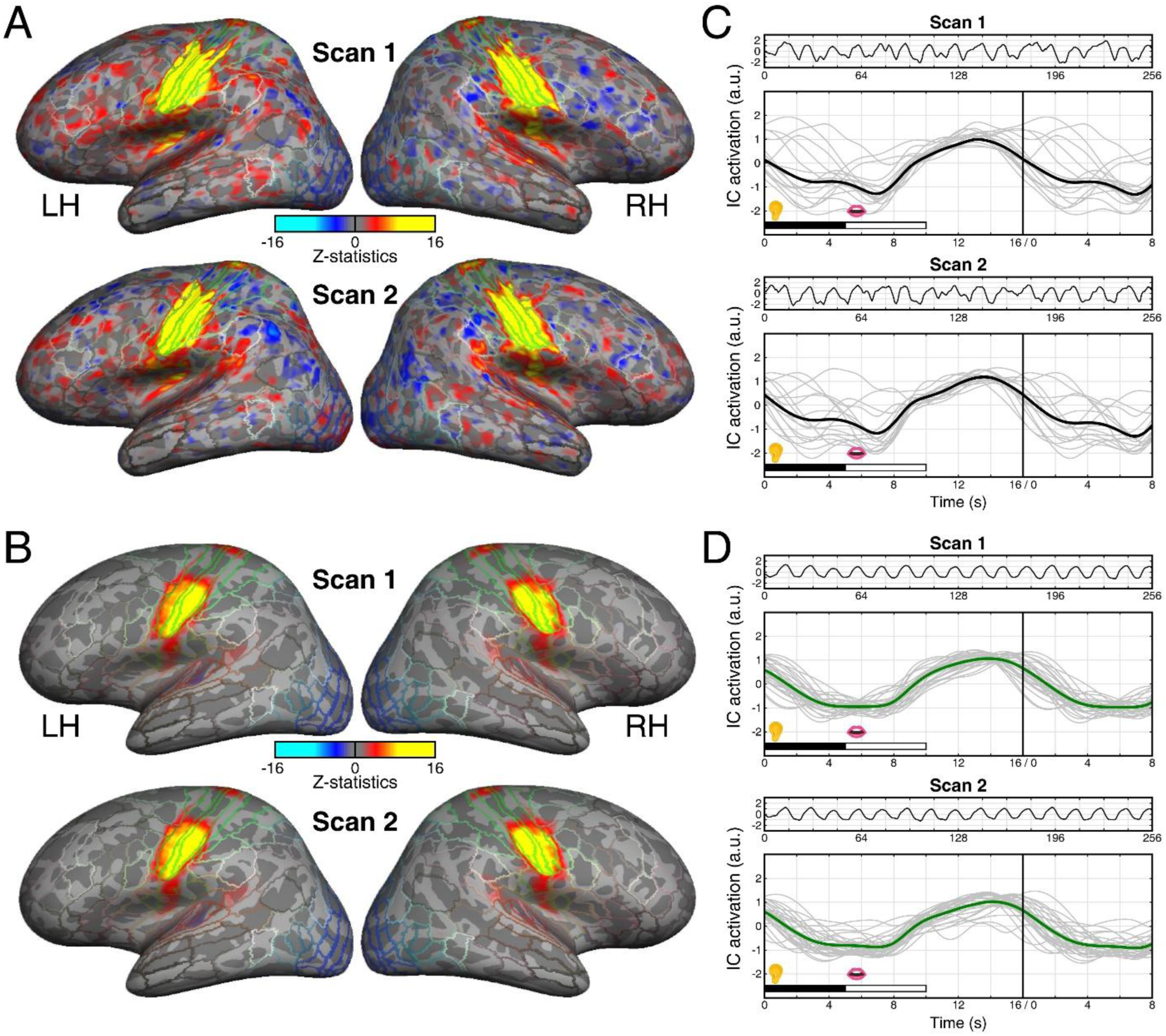
Spatial maps and time courses of the SMC IC in the listening-reciting task. (A) Spatial maps of the SMC IC from two scans of a single subject. (B) Group-average (*N* = 31) spatial maps of the SMC IC from two scans. (C) Time courses associated with the SMC IC in (A). The first and third panels show the full 256-s time courses for both scans. The second and fourth panels show the time courses in 24-s epochs (gray curves) and their averages (black curves) within each scan. (D) Time courses associated with the SMC IC in (B). The first and third panels show the full 256-s time courses averaged across 31 and 29 subjects for Scan 1 and Scan 2, respectively. The IC time courses from two subjects in Scan 2 (the same as those excluded in Fig. 4) were excluded from group averaging and subsequent analysis. The second and fourth panels show the time courses in 24-s epochs averaged within each subject (gray curves; *N* = 31 for Scans 1 and 2) and across subjects (red curves; *N* = 31 for Scan 1 and *N* = 29 for Scan 2). All conventions are consistent with those in Fig. 3.

**Fig. 7.**
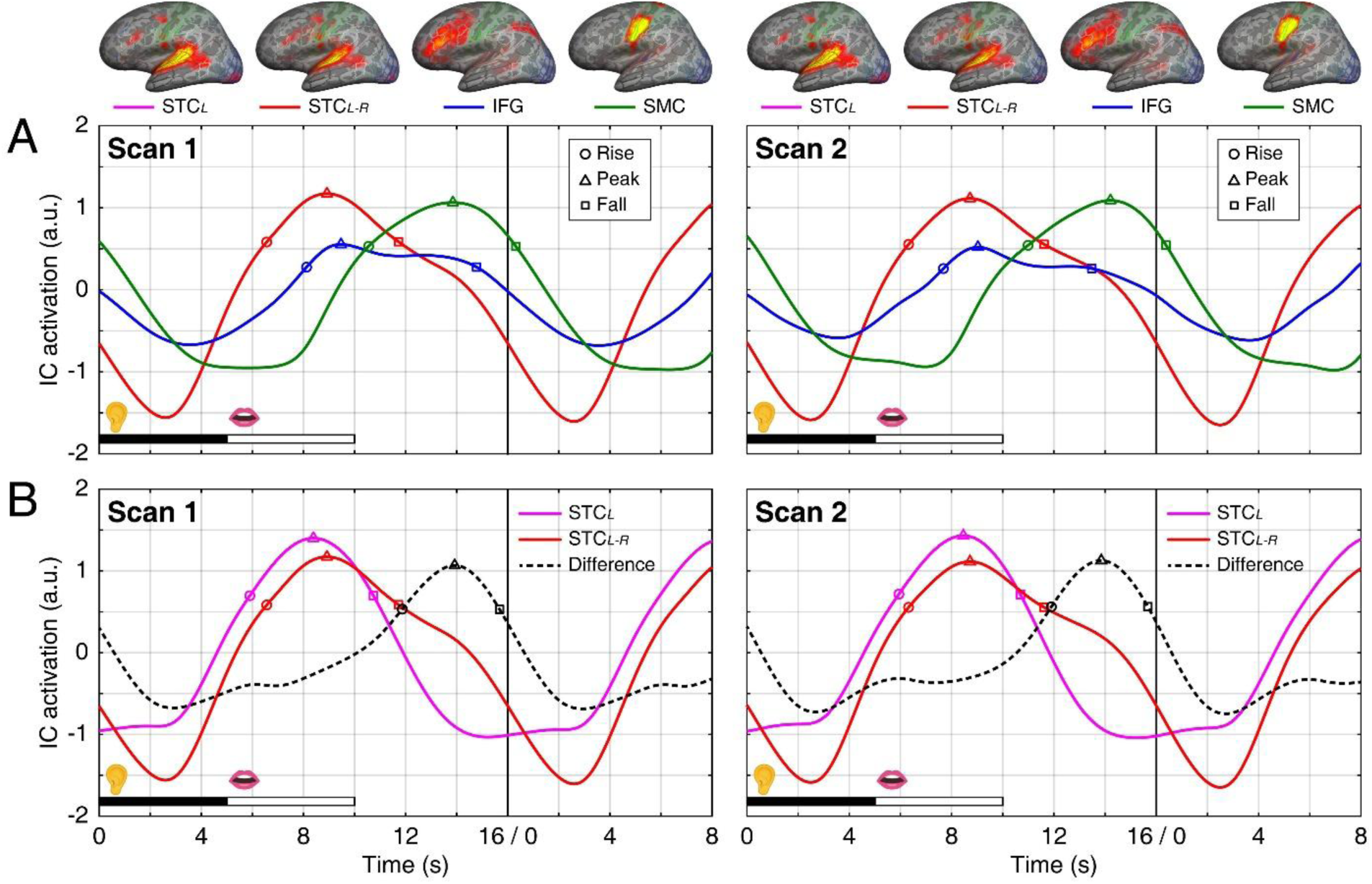
Group-average IC activation time courses. (A) Overlay of group-average time courses of the ICs identified for the listening-reciting task (redrawn from Figs. 4-6). (B) Each black dashed curve indicates the difference between the time courses of the STC*_L-R_* IC (listening-reciting task, red curve; redrawn from Fig. 4) and STC*_L_* IC (listening task, magenta curve; redrawn from Fig. 3) tasks. The top row shows the group-average IC spatial maps in the left hemisphere (redisplayed from Figs. 3-6) for Scan 1 and Scan 2, respectively. The color coding for group-average IC time courses is consistent with that in Figs. 3-6. The circle, triangle, and square indicate the timings of the rise, peak, and fall in the group-average hemodynamic response profile. Exact values are provided in Table 1.

**Table 1.** Key timings of hemodynamic responses in group-average IC time courses.

|  | Task | Listening | Listening-reciting |  |  | Self-monitoring |
| --- | --- | --- | --- | --- | --- | --- |
| | IC | $STC_L$ | $STC_{L-R}$ | IFG | SMC | $STC_{L-R} - STC_L$ |
| Scan 1 | Rise | 5.90 | 6.56 | 8.12 | 10.55 | 11.87 |
|  | Peak | 8.39 | 8.92 | 9.47 | 13.85 | 13.92 |
|  | Fall | 10.74 | 11.72 | 14.78 | 16.32 | 15.68 |
| Scan 2 | Rise | 5.94 | 6.32 | 7.68 | 10.99 | 11.90 |
|  | Peak | 8.46 | 8.72 | 9.03 | 14.21 | 13.85 |
|  | Fall | 10.69 | 11.61 | 13.48 | 16.38 | 15.67 |

All values were measured from the trial onset and are expressed in seconds.

For each IC time course, the timings peak (maximum point at 𝑡^′^, measured from the zero baseline), rise (ascending half-maximum point at 𝑡^′^*_p_*), and fall (descending half-maximum point at 𝑡^′^*_r_*) as well as the activation duration (full width at half maximum; FWHM = 𝑡^′^*_f_*− 𝑡^′^*_r_*) were identified from the hemodynamic response in each within-subject-average epoch, 𝑋^′^*_S_*(𝑡^′^*_e_*), and each group-average epoch, 𝑋^′^*_G_*(𝑡^′^*_e_*) (Figs. 7, 8).

## 3. Results

### 3.1 Head motion analysis

Our comprehensive procedures and meticulous setup for fMRI experiments (Fig. 1) successfully constrained the range (maximum - minimum) of head motion to less than 1 mm in translation and 1° in rotation within each scan across tasks and subjects (Fig. 2). The efficacy of training and head-constraining measures was particularly evident in the displacements observed in the left-right (median range < 0.1 mm) and posterior-anterior (median range < 0.25 mm) directions, as well as the minimal rotation in roll and yaw (median range < 0.2°). However, minor head motions were observed in the superior-inferior direction (median range < 0.4 mm) and in pitch (median range < 0.4°), which were slightly more pronounced during the listening-reciting task.

The high consistency of these motion mitigation strategies across subjects was confirmed by the small variations (standard deviations) in motion parameters, which were less than 0.1 mm in translation and 0.1° in rotation in most scans (Fig. 2).

### 3.2 Independent brain processes

**Speech perception (listening task).** Across 31 subjects, independent component analysis on each functional scan of the listening task resulted in 78.2 ± 8.7 and 82 ± 9.3 ICs for Scan 1 and Scan 2, respectively. In a single subject, an IC exhibiting speech perception-related activations predominantly in the superior temporal cortex (STC) was identified among all ICs decomposed from each scan (Fig. 3A). The STC IC map shows activations spanning the posterior lateral sulcus, superior temporal gyrus (STG), and the middle to posterior portions of the superior temporal sulcus (mid-post STS) in both hemispheres. Furthermore, small activation clusters were found in the premotor cortex, inferior frontal gyrus (IFG), and parietal operculum. Surface-based group averaging of the STC IC maps revealed an activation pattern consistently observed across subjects, with strong activation in a region extending from the posterior lateral sulcus to STG (Fig. 3B). These areas include TA2, 52, MBelt, A1, LBelt, PBelt, A4, A5, Rl, and PSL in the HCP-MMP1 atlas (Glasser et al., 2016). Additionally, areas 55b, IFJa, IFSp, IFSa, OP4, STV, TPOJ1, TPOJ2, PHT, STSvp, and STSdp showed weaker activations. The STC IC time courses consistently exhibited periodic BOLD signals (16 cycles/scan) in response to external speech in both scans within and across subjects (first and third panels in Fig. 3C, D). Within the 16-s trial cycle, the average time courses displayed a roughly bell-shaped temporal profile (second and fourth panels in Fig. 3C, D).

**Speech perception and production (listening-reciting task).** Across 31 subjects, independent component analysis on each functional scan of the listening-reciting task resulted in 101.5 ± 18.7 and 103 ± 17.9 ICs for Scan 1 and Scan 2, respectively. Three ICs associated with the speech perception, planning, and production processes were identified in both scans within and across subjects (Figs. 4-6). The first IC (STC) exhibited an activation pattern similar to the STC IC identified for the listening task (Fig. 4A, B). The second IC (IFG) exhibited strong activations in the inferior frontal gyrus, as well as weaker activations in the ventral premotor cortex, dorsal lateral prefrontal cortex, and the posterior parietal cortex (Fig. 5). These areas include p47r, IFSa, IFSp, IFJa, p9-46v, IFJp, 6r, PEF, AVI, IP1, IP2, and LIPd in the HCP-MMP1 atlas. The third IC exhibited strong activations predominantly in the orofacial representations of the sensorimotor cortex (SMC, areas 4, 3a, 3b and 1; Sereno & Huang, 2006; Huang et al., 2012; Huang & Sereno, 2018), as well as weaker activations in areas OP4 and OP2-3 in the HCP-MMP1 atlas (Fig. 6). The areas in the operculum correspond to an auditory area, 43aud, and the secondary somatosensory cortex (S-II) in the CsurfMap1 atlas (Sereno et al., 2022).

The STC IC time courses exhibited periodic BOLD signals (16 cycles/scan) in response to external and self-generated speech in both scans within and across subjects (first and third panels in Fig. 4C, D). Compared with STC, the IFG IC time courses exhibited delayed activations with lower amplitudes, reflecting the planning process for speech production. Finally, the SMC IC time courses exhibited the latest activations induced during speech production. The relative timings of activations among these ICs are further examined in the following section.

### 3.3 Time-resolved hemodynamic responses

Despite substantial overlap, the group-average hemodynamic response profiles of STC, IFG, and SMC ICs exhibited temporal shifts reflecting sequential brain processes during the listening-reciting task (Fig. 7A). The timings of the rise, peak, and fall of the average hemodynamic response are summarized for each IC in Table 1. While spatial ICA revealed different ICs associated with speech perception, planning, and production processes, we did not identify separate ICs for processing external speech and self-generated speech in the listening-reciting task. Instead, we identified a single IC (STC*_L-R_*) with a spatial map similar to the STC*_L_* IC identified for the listening task. Compared with the STC*_L_*, the STC*_L-R_* IC exhibited a widened and delayed activation profile. According to the superposition principle of linear systems (Boynton et al., 2012), we assumed that the widened profile of the STC*_L-R_* IC resulted from a linear combination of hemodynamic responses to external speech and self-generated speech. The time course of the STC*_L_* IC, serving as a reference for its response to external speech, was then subtracted from the time course of the STC*_L-R_* IC. The estimated hemodynamic response (black dashed curve in Fig. 7B) peaked at 13.92 s for Scan 1 and 13.85 s for Scan 2, which closely aligned with the activation peak of the SMC IC (13.85 s for Scan 1 and 14.21 s for Scan 2; Table 1, Fig. 7A). The result suggested that the time course of the STC*_L-R_* IC consisted of a first hemodynamic response to external speech during the perception phase (0–5 s) and a second hemodynamic response to self-generated speech during the reciting phase (5–10 s).

### 3.4 Temporal relationships between speech production and hemodynamic responses

The timings of speech onset and offset, together with the rise, peak, and fall of hemodynamic responses were examined across subjects for three ICs identified for the listening-reciting task (Fig. 8; Table 2). In addition to the median and interquartile range (IQR), the mean and standard deviation of the peak timing of each IC’s hemodynamic response were compared with the onset of speech output (6.15 ± 0.34 s and 5.98 ± 0.28 s for Scan 1 and Scan 2, respectively).

**Fig. 8.**
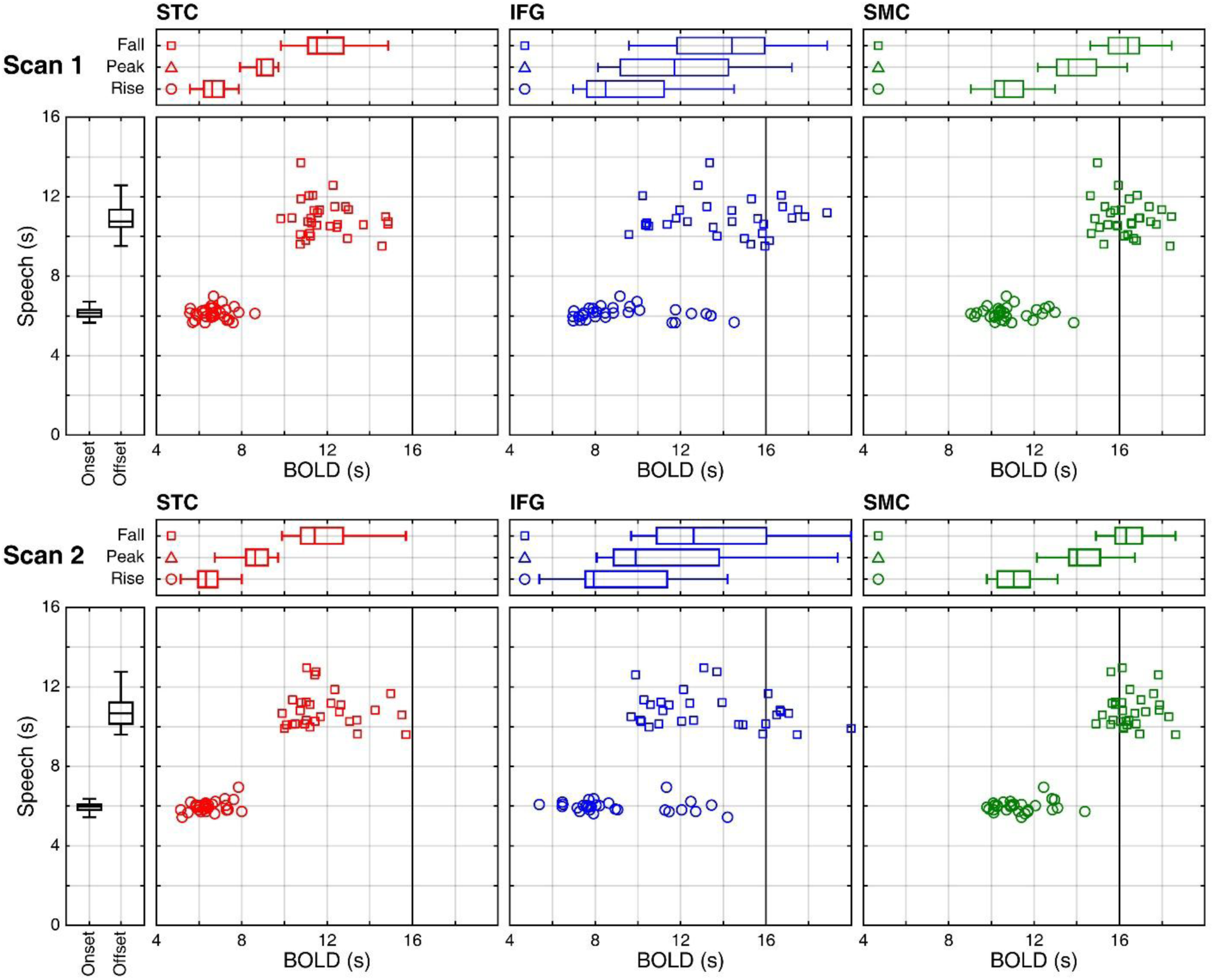
Distributions of key timings of speech output and hemodynamic responses in the listening-reciting task. For each scan, the horizontal panels display the distributions of the rise, peak, and fall times of the hemodynamic responses (BOLD) in the STC (red), IFG (blue), and SMC (green) ICs. The vertical panels show the distributions of speech onset and offset times. *N* = 31 for Scan 1 and *N* = 29 for Scan 2. In the scatter plots, each marker indicates a subject’s data point. Open circles: joint distribution of the speech onset and hemodynamic response rise across subjects. Open squares: joint distribution of the speech offset and hemodynamic response fall across subjects.

**Table 2.** Median and interquartile range (IQR) of key timings of speech output and hemodynamic responses in IC time courses.

|  | Speech |  | Hemodynamic response |  |  |  |
| --- | --- | --- | --- | --- | --- | --- |
|  | Onset | Offset | IC | STC | IFG | SMC |
| Scan 1 | 6.15 (0.34) | 10.75 (0.87) | Rise | 6.61 (0.93) | 8.48 (3.62) | 10.59 (1.35) |
|  |  |  | Peak | 8.92 (0.77) | 11.72 (5.06) | 13.61 (1.87) |
|  |  |  | Fall | 11.52 (1.66) | 14.4 (4.1) | 16.39 (1.45) |
| Scan 2 | 5.98 (0.28) | 10.67 (1.08) | Rise | 6.32 (0.92) | 7.93 (3.83) | 11.05 (1.53) |
|  |  |  | Peak | 8.63 (1.02) | 9.89 (4.93) | 14.01 (1.45) |
|  |  |  | Fall | 11.42 (1.98) | 12.61 (5.13) | 16.31 (1.26) |

All values were measured from the trial onset and are expressed in seconds. $N = 31$ for Scan 1; $N = 29$ for Scan 2.

For the STC IC, the mean times of hemodynamic response peak were 9.21 ± 1.2 s (Scan 1) and 8.85 ± 1.13 s (Scan 2) from trial onset, with average lags of 3.06 s and 2.87 s after the onset of speech output for Scan 1 and Scan 2, respectively. The mean times for the IFG IC to reach peak from trial onset occurred later at 11.64 ± 2.68 s (Scan 1) and 11.31 ± 3.01 s (Scan 2), with average lags of 5.49 s and 5.33 s after the speech onset for Scan 1 and Scan 2, respectively. The hemodynamic response of the SMC IC peaked at 14.06 ± 1.11 s (Scan 1) and 14.3 ± 1.16 s (Scan 2) from trial onset, with the longest post-speech-onset lags of 7.91 s and 8.32 s for Scan 1 and Scan 2, respectively.

The temporal relationships between speech onset and the rise of the hemodynamic responses in the STC IC display the most tightly lustered distributions in both scans, followed by the SMC IC with slightly wider distributions, and the IFG IC with highly dispersed distributions (open circles in Fig. 8, see also IQR values in Table 2). The temporal relationships between speech offset and the fall of the hemodynamic responses in the SMC IC display the most concentrated distributions in both scans, while the STC and IFG ICs show dispersed distributions (open squares in Fig. 8; see also IQR values in Table 2).

Overall, ICs in the auditory and sensorimotor cortices, compared with cognitive processing regions in the frontal cortex, were more predictable due to the fixed timings of the stimulus phase (0-5 s) and production phase (5-10 s) within each trial (see Discussion).

## 4. Discussion

Since the 1990s, fMRI has been one of the noninvasive neuroimaging techniques used for investigating the neural basis of cognition. Although fMRI offers high spatial resolution for mapping brain functions, the rapid switching of gradient magnetic fields during echo planar imaging generates loud noise, posing a major challenge for fMRI investigations of the auditory system and speech processing. For an experiment to successfully detect the neural response to a delivered auditory stimulus, that response must be sufficiently robust to rise above the competing neural activity evoked by the scanner noise itself (Dewey, et al., 2021).

To mitigate the interference due to scanner noise, sparse-sampling fMRI designs have been developed and used study speech processing (Erb et al., 2013; Golfinopoulou et al., 2011; Hervais-Adelman et al., 2015; Longcamp et al., 2019; Murray et al., 2022; Orellana et al., 2014; Perrachione & Ghosh, 2013; Tao et al., 2026). While effective at creating quiet periods for stimulus presentation and over speech production, this approach imposes significant constraints on experimental designs and analyses. Prolonged trial durations are required to accommodate both the stimulus presentation and the subsequent lag of the hemodynamic response, all while ensuring sufficient data is collected for statistical power. For instance, in a sound mimicry study that integrated both perception and production within a single trial, sparse-sampling designs with an interval of 10 s would require a long scan time to obtain sufficient trials for statistical analysis (Lewis et al., 2018). Furthermore, the significant data loss due to sparse sampling makes it unable to characterize the full temporal profile of brain dynamics during complex, multi-stage tasks like auditory perception followed by overt production.

In addition to acoustic noise, head motion artifact remains a significant source of variance in fMRI data, capable of introducing spurious correlations and obscuring genuine neural signals (Liu, 2016). In the present study, we implemented a comprehensive strategy to minimize head motion throughout the functional session, allowing tasks involving overt production of 5-s sentences. While the efficacy of individual methods like custom-molded headcases or masks can be debated (Jolly et al., 2020; Power et al., 2019), our approach combined several practices: (1) subjects underwent a training session with real-time head motion feedback to train them to remain still in a mock scanner; (2) subjects wore custom-molded thermoplastic masks for rigid head stabilization; and (3) the head motion was further confined by filling residual air gaps within the head coil. This concerted effort effectively restricted the range of head displacement within 1 mm and rotation below 1° per scan in all participants (Fig. 2), providing a stable foundation for the high-resolution analyses to follow.

By overcoming the challenges associated with speech perception and overt production in the MRI scanner, we achieved continuous sampling with real-time auditory feedback and minimal head-motion artifacts. These technical solutions now allow us to address advanced questions in speech processing that were previously inaccessible with sparse-sampling designs. In an event-related fMRI experiment, as long as two auditory stimuli (events) are separated by a prolonged interval, a peak of hemodynamic response to each stimulus can be distinctly identified from the BOLD time course (Hickok et al., 2003). However, the hemodynamic responses to two stimuli would not be easily separated when they are presented in close succession.

In this study, we explored the use of ICA for decomposing independent brain processes associated with sentence-level speech perception and production. Each functional scan was subjected to ICA and the resulting IC maps were displayed on the cortical surface reconstructed from the structural images of each individual subject. We then identified IC maps with similar spatiotemporal activation patterns across subjects and combined them through surface-to-surface registration to obtain surface-based group-average IC maps (Fischl et al., 1999b; Sereno et al., 2022). This subject-level ICA approach is fundamentally different from the volume-based group-ICA method (e.g., Janssen & Mendieta, 2020), where the functional volumes were combined across subjects before being subjected to ICA. Our method yields subject-level IC maps and time courses with high spatial and temporal precision, enabling accurate functional localization and detection of intersubject variations. As shown in panel D across Figs. 3-6, the overall waveforms of IC time courses are consistent across subjects, while the key timings of hemodynamic responses vary (Fig. 8).

Across subjects, we identified an IC in the superior temporal cortex, an IC in the inferior frontal gyrus, and an IC in the orofacial sensorimotor cortex. The time courses of these ICs exhibited sequential activations corresponding to the speech perception (STC), articulatory planning (IFG) (Basilakos et al., 2018), and articulation (SMC) phases during the listening-reciting task (Figs. 4-6). While the spatial ICA method used in the current study did not resolve separate ICs associated with the processing of external speech and self-generated speech, it revealed that the similar STC ICs identified for the listening and listening-reciting tasks exhibited different activation profiles. Compared with the listening task, the listening-reciting task induced a wider profile of hemodynamic response in the STC IC (Fig. 7B). With a sampling rate of 1 s, we were able to upsample and contrast the IC time courses between listening-reciting and listening tasks. Based on the properties of a linear system (Boynton et al., 2012), we postulated that the widened profile resulted from the linear superposition of two hemodynamic responses – one to external speech during the perception phase (0-5 s) and the second to self-generated speech during the production phase (5-10 s). Subtracting the STC IC time course of the listening task from that of the listening-reciting task indeed revealed a second hemodynamic response (Fig. 7B), with its peak timing aligned to that of the SMC IC. In some subjects, the second hemodynamic response to self-generated speech appeared as a small hump embedded in the descending section of the widened profile in the STC IC (Fig.4D).

In addition to ICA decomposition, continuous sampling enables the examination of the temporal relationships between sentence-level speech production and the evoked hemodynamic responses with high precision. The hemodynamic profiles of the STC (listening task) and the SMC represent the BOLD responses to the onset of stimulus and onset of speech output respectively. For the listening task, the peak of the STC IC time course occurred ∼8.5 s after the onset of a spoken sentence (external speech) (Fig. 7B, Table 1). For the listening-reciting task, the peak of the SMC IC time course occurred ∼8 s after the onset of speech output at ∼6 s within a 16-s trial (Fig. 7A, Table 2). These peak latencies, later than the 5 to 6 s range in typical hemodynamic responses, indicated extended processing time for sentence-level stimuli. Furthermore, the peak latencies in the STC and SMC ICs were less dispersed than those in the IFG IC, suggesting stable hemodynamic responses in the auditory and sensorimotor cortices across subjects due to the well-defined timings of stimulus and response phases.

The hemodynamic response profiles of the IFG IC may represent the speech rehearsing and articulatory planning processes and do not directly align to the timing of a stimulus. Indeed, the IFG IC time courses exhibited large inter-subject variability in the duration between the speech onset and hemodynamic peak (Fig. 8, Table 2), suggesting differential strategies in rehearsal (e.g. some subjects may start articulatory planning as soon as the onset of stimulus while some may only start after a whole sentence was heard). This variability underscores a critical limitation of sparse-sampling designs. As the typical window for image acquisition is limited to 2 s per trial (Hervais-Adelman et al., 2015; Perrachione & Ghosh, 2013; Tao et al., 2026), sparse-sampling fMRI could not reliably capture the temporally variable responses to high-level language processing in frontal regions. Future sparse-sampling designs involving sentence-level stimuli could therefore incorporate longer and more flexible acquisition windows, informed by the temporal profiles revealed through continuous sampling approaches.

While informative, this study entails some limitations. First, our listening-reciting task was designed to enable precise control over speech onset and provide well-defined perception and production phases. The listening-reciting task does not fully capture the cognitive demands of natural, generative speech production. Rather, it relies predominantly on the phonological loop component of working memory (Baddeley, 2003). Consequently, the neural dynamics we observed, particularly in the inferior frontal gyrus, may reflect the articulatory planning of rehearsal to a greater extent than a higher-level linguistic formulation. Nonetheless, the controlled nature of this task served as a critical first step for validating the feasibility of continuous sampling and in disentangling temporally overlapping hemodynamic responses.

Second, the spatial ICA method employed here did not yield separate independent components associated with the processing of external speech versus self-generated speech. This likely reflects the substantial spatial overlap of neural populations within the superior temporal cortex that subserve both auditory perception of external speech and self-monitoring of one’s own vocal output (Tourville & Guenther, 2011). Such overlapping functional networks may not be resolvable at the spatial and temporal resolution of conventional fMRI, highlighting the need for future studies to employ complementary techniques such as multivoxel spatiotemporal pattern analysis to more effectively dissociate these mixed signals.

In conclusion, we present methodological advancements that allow continuous fMRI sampling during sentence-level speech perception and production tasks. Independent component analysis revealed time-resolved hemodynamic responses across multiple ICs associated with the perception, planning, production, and self-monitoring processes. The temporal information identified, such as the delay between speech onset and hemodynamic peak, may facilitate future auditory fMRI studies in determining the optimal timings for stimulus presentation, overt production, and image acquisition for either continuous- or sparse-sampling designs. This study not only demonstrates the feasibility of naturalistic speech tasks but also lays the groundwork for future research into tasks like simultaneous interpreting and collaborative singing, and explores the nuanced interactions between externally perceived auditory input and awareness of internally generated sounds, advancing our knowledge of cognitive processing in these multifaceted activities.

## Data and code availability

The data that support the findings of this study are available from the corresponding author upon request. Custom codes for analyzing fMRI data are included in *csurf* (a FreeSurfer-compatible package) available at https://pages.ucsd.edu/~msereno/csurf/

## Author contributions

T.I.L., B.L.R., D.L., V.L.C.L., and R.S.H. designed the research. T.I.L., C.T.L., C.U.C., and R.S.H. collected the data. T.I.L., A.L., J.H.A., C.U.C., M.I.S., and R.S.H. analyzed the data. T.I.L., V.L.C.L., and R.S.H. wrote the paper. B.L.R., D.L., V.L.C.L., and R.S.H. supervised the work.

## Acknowledgments

We thank Yi Tang and Mengying Guo for help with fMRI experiments; Yafang Li and Ut Meng Lei for data preprocessing.

